# *Klebsiella* MALDI TypeR: a web-based tool for *Klebsiella* identification based on MALDI-TOF mass spectrometry

**DOI:** 10.1101/2020.10.13.337162

**Authors:** Sébastien Bridel, Stephen C. Watts, Louise M. Judd, Taylor Harshegyi, Virginie Passet, Carla Rodrigues, Kathryn E. Holt, Sylvain Brisse

**Affiliations:** Biodiversity and Epidemiology of Bacterial Pathogens, Institut Pasteur, Paris, France; Department of Infectious Diseases, Central Clinical School, Monash University, Melbourne, Victoria 3004, Australia; London School of Hygiene & Tropical Medicine, London WC1E 7HT, UK

## Abstract

**Motivation:** *Klebsiella* species are increasingly multidrug resistant pathogens affecting human and animal health and are widely distributed in the environment. Among these, the *Klebsiella pneumoniae* species complex (KpSC), which includes seven phylogroups, is an important cause of community and hospital infections. In addition, the *Klebsiella oxytoca* species complex (KoSC) also causes hospital infections and antibiotic-associated haemorrhagic colitis. The unsuitability of widely used clinical microbiology methods to distinguish species within each of these species complexes leads to high rates of misidentifications that are masking the true clinical significance and potential epidemiological specificities of individual species.

**Results:** We developed a web-based tool, *Klebsiella* MALDI TypeR, a platform-independent and user-friendly application that enables uploading raw data from MALDI-TOF mass spectrometer to identify *Klebsiella* isolates at the species complex and phylogroup levels. The tool is based on a database of previously identified biomarkers that are specific for either the species complex, individual phylogroups, or related phylogroups, and is available at https://maldityper.pasteur.fr.

## 1 Introduction

Recent works pointed out the difficulties that microbiology laboratories encounter in the identification of individual members of the *K. pneumoniae* and *K. oxytoca* species complexes (Brisse *et al.*, 2004; Seki *et al.*, 2013; Fonseca *et al.*, 2017; Long *et al.*, 2017). Indeed, each species complex encompasses several closely related phylogroups, which correspond to species or subspecies with few discriminatory markers (Rodrigues *et al.*, 2018; Merla *et al.*, 2019). As a consequence, the true clinical significance of these phylogroups and their potential epidemiological specificities are poorly defined. The current Bruker Biotyper Matrix Assisted Laser Desorption Ionization - Time of Flight (MALDI-TOF) mass spectrometry (MS) database comprises reference spectra only for *K. pneumoniae*, *K. variicola* and *K. oxytoca* (Berry *et al.*, 2015; Long *et al.*, 2017; Dinkelacker *et al.*, 2018). However, recent work demonstrated the ability of MALDI-TOF MS to generate discriminant peaks at phylogroup level (Rodrigues *et al.*, 2018; Merla *et al.*, 2019; Dinkelacker *et al.*, 2018). Here, we designed *Klebsiella* MALDI TypeR in order to leverage these biomarkers for improved accuracy and resolution of *Klebsiella* species identification from MALDI-TOF data.

## 2 Features

*Klebsiella* MALDI TypeR is an application with a web interface designed with R and R Shiny. A user can upload either a single spectrum or multiple spectra (a maximum number of spectra is not defined, but we recommend working with batches of 100 spectra per analysis). The uploaded spectra must be in zip archive format. Each archive may correspond either to a single raw spectrum or several raw spectra of the same isolate (technical replicates). For better results, we recommend using at least 3 technical replicates for each isolate. The spectra shape can be checked after their upload: a panel shows each uploaded spectrum for quick visual inspection (intensity checking) and a table highlights the presence (if any) of empty spectra. For identification, the tool is based on a 2-step process. The first step differentiates isolates at the species complex level using two sets of biomarkers, each set corresponding to reference biomarkers for (i) the *Klebsiella pneumoniae* species complex (29 biomarkers, Table S1) or (ii) the *Klebsiella oxytoca* species complex (57 biomarkers, Table S1). This step can be performed, based on user choice, with one of two alternative algorithms for peak detection. The first one [using the R package MALDIquant (Gibb and Strimmer, 2012)] is set by default and runs faster. The second one [using the MassSpecWavelet package (Du *et al.*, 2006)] generally detects more significant peaks but is slower. Species complex identification is based on the ratio of common peaks between the sample peak list and a reference peak list, to the number of peaks in the corresponding reference (this ratio is called “Identification Value”). The input sample is assigned to the species complex with the highest Identification Value. The second step is to identify the specific phylogroup (corresponding to a species or subspecies) within that complex. The raw spectrum is processed again by the method known as Multi Peak Shift Typing (MPST) (Bridel *et al.*, 2020), but this time using the set of biomarkers corresponding to the assigned species complex, in order to recalibrate *m/z* values. The biomarkers used for the two MPST schemes (one for each species complex, see Tables S2 to S5) are based on previous work (Rodrigues *et al.*, 2018; Merla *et al.*, 2019). Briefly, an MPST scheme is analogous to a multi-locus sequence typing (MLST) scheme where an allele type (AT) corresponds to an isoform (IF) for one protein (each IF is defined by a specific mass – or a *m/z* value on the spectrum) and a MALDI-Type (MT) corresponds to a sequence type (ST). Each MT was observed within a single phylogroup. The tool attempts to define the IF for each biomarker and to attribute the spectrum to the phylogroup corresponding to the relevant MT. The only exception to this process was introduced for the discrimination of *K. oxytoca* phylogroups Ko1, Ko4 and Ko6 (*K. oxytoca, K. pasteurii* and *K. grimontii*, respectively). These three species form a phylogenetic branch of closely related species within the *K. oxytoca* species complex (Merla *et al*., 2019), and their spectra are very similar such that they share the same MT. In order to distinguish among these three phylogroups, we added 3 previously described biomarkers (Merla *et al.*, 2019) for which the tool checks for presence/absence within the spectra (i.e. these 3 biomarkers do not correspond to peak shifts). If any of the above biomarkers (peak shifts or presence/absence) are not found, automatic identification will not be possible, and the result is reported to the user as NT (not-typable). In this case, manual analysis or data re-acquisition are needed. To help resolve these cases and allow users to further explore the data, we included a feature to interactively explore the spectra. The plot window shows the sample spectra overlaid on the reference spectra (1 to 3 spectra for each phylogroup). Predefined windows (one for each biomarker) allow the user to directly investigate specific spectral regions. This feature could also enable the detection of new isoforms or biomarkers. It also contains a customizable option that allows the user to select any region of the spectra. Examples of these features are depicted in Figures S1 to S3.

Website is available at http://hub05.hosting.pasteur.fr/sbridel/klebsiellatyper/ (or https://maldityper.pasteur.fr

## 3 Validation

We first tested fifty *K. pneumoniae* species complex (KpSC) isolates grown overnight on lysogeny broth (LB) agar (37°C, 18h) and subjected to ethanol/formic acid extraction following the manufacturer recommendations (Bruker Daltonics, Bremen, Germany). For each strain, a total of 3 spectra from the same spot were acquired (Bruker Microflex LT mass spectrometer, Paris). This dataset yielded correct assignments to species complex for 50/50 (100%) isolates, 47 (94%) of which had an identification value >50% (Table S6 and S7). The same samples prepared using different experimental conditions (two media: LB and Columbia agar plus 5% sheep blood; each with both extraction and direct application in the MALDI-TOF target) also yielded correct complex assignments for all isolates, albeit with lower confidence (Table S1). Regarding phylogroup assignment, using LB-extraction 90% of samples were correctly identified at phylogroup level (Table S7), while the other sample preparation methods yielded better results overall (Table S8). To validate robustness of the tool to instrument and user variation, we tested an additional dataset comprising 248 KpSC isolates prepared using the LB-extraction method (176 isolates with a single spectra, 26 in duplicate and 68 in triplicate); these spectra were generated on a different instrument (Bruker MALDI Biotyper microflex LT-SH) in a different laboratory (Melbourne, Australia). 244/248 isolates (98%) were assigned to the correct species complex, and 219/244 (90%) were correctly identified at the phylogroup level (84% to 100% for each phylogroup, see Table S9).

*K. oxytoca* species complex (KoSC) assignments were tested using 28 isolates prepared using LB-extraction and triplicate spectra generated on the Bruker Microflex LT mass spectrometer instrument (Paris). All isolates were correctly assigned to complex level and most (89%) were assigned to the correct MALDI-Type (Tables S10 and S11). Discrimination of MALDI-Type MT1 (n=15, 54%) into phylogroups Ko1, Ko4 and Ko6 was poor (one isolate correctly assigned as Ko1) because the three additional biomarkers intended to discriminate between these phylogroups were captured only rarely (Table S6).

## 4 Conclusions

We developed the *Klebsiella* MALDI TypeR web tool for identification of *Klebsiella* species based on MALDI-TOF MS. Our tool allows the fast analysis of single or multiple spectra, including their visualization. We show that MALDI-TOF MS leads to accurate identification for *Klebsiella* phylogroups of the KpSC, the major *Klebsiella* species complex found in clinical samples. *Klebsiella* MALDI TypeR therefore represents an important resource for the identification of the KpSC members, which are currently difficult to identify in routine microbiology practice. For the KoSC, we found promising data, but further validation based on a larger dataset would be required. The biomarker approach may not be suitable for accurate discrimination of Ko1, Ko4 and Ko6. In the future, the website may be updated to integrate additional methods for identification and typing, such as supervised learning.

## Acknowledgements

This work was financially supported by the MedVetKlebs project, a component of European Joint Programme One Health EP, which has received funding from the European Union’s Horizon 2020 Research and Innovation Programme under Grant Agreement No. 773830. This work was also supported by the kind help of Stevenn Volant and Amine Ghozlane from Bioinformatics and Biostatistics HUB (R Shiny tips). The web tool is also hosted by the team Bioinformatics and Biostatistics HUB at Institut Pasteur.

## Notes

http://hub05.hosting.pasteur.fr/sbridel/klebsiellatyper/

### Competing Interest Statement

The authors have declared no competing interest.

