## Supplemental Figure S1 for "*Klebsiella* MALDI TypeR: a web-based tool for *Klebsiella* identification based on MALDI-TOF mass spectrometry"

### A – Load and check spectra

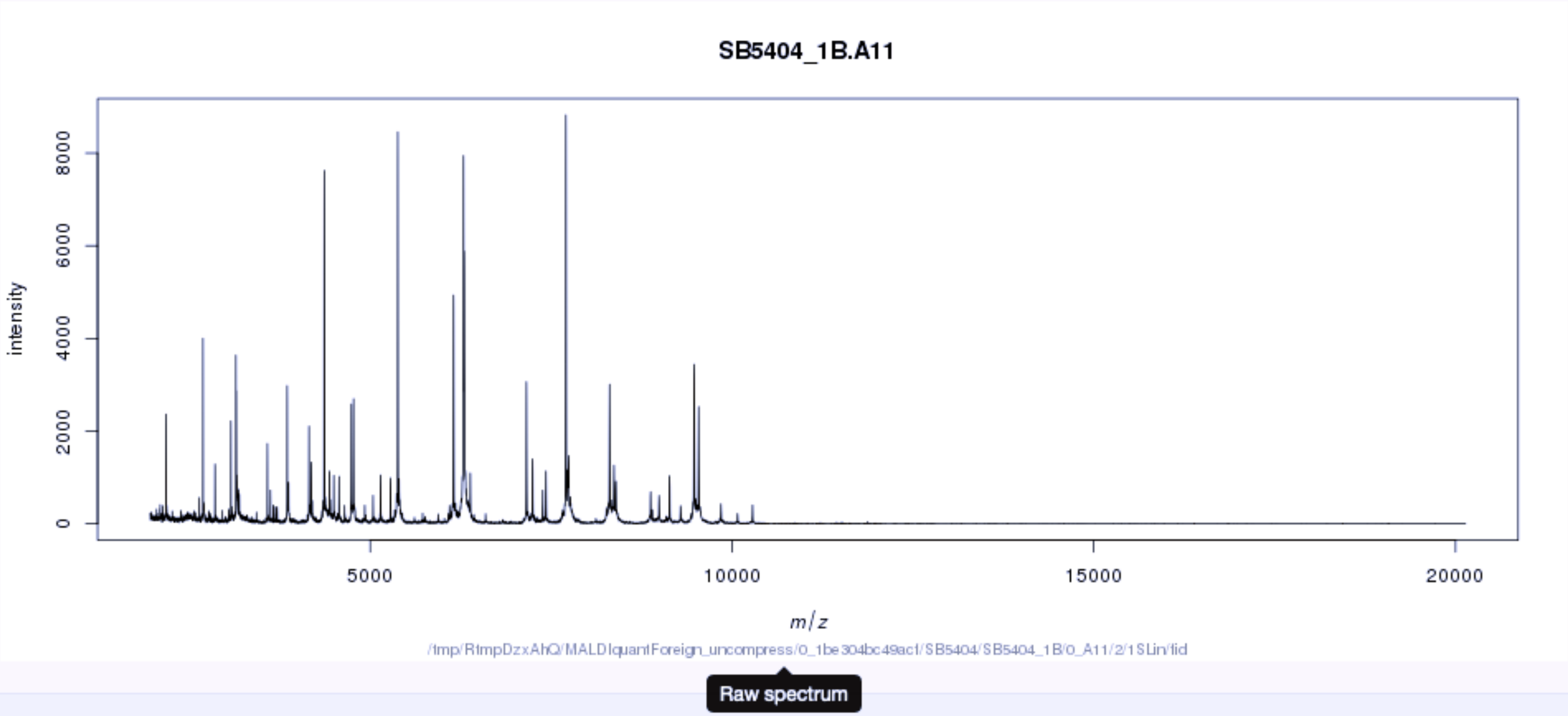


Show 10 entries

Search:

| Isolate File | Loaded Spectra | Total Spectra | Index Range | Status |
|---|---|---|---|---|
| SB5404.zip | 3 | 3 | 1-3 | All spectra loaded |

Showing 1 to 1 of 1 entries

Previous 1 Next
