## Supplemental Figure S2 for "*Klebsiella* MALDI TypeR: a web-based tool for *Klebsiella* identification based on MALDI-TOF mass spectrometry"

### B – Two steps identification

#### B1 – Species complex identification – Summary table

| Sample | Detected Peaks | Proposed identification | Common Peaks | Reference peaks | Similarity ratio | Identification Strength |
|---|---|---|---|---|---|---|
| SB5404 | 72 | *K. pneumoniae species complex* | 16 | 29 | 55.17 | good |

#### B2 – Phylogroup identification – Summary table

| Sample | Proposed species identification |
|---|---|
| SB5404 | *K. pneumoniae* |

#### B3 – Phylogroup identification – Detailed table (biomarkers table)

| sample | YdgH** | YdgH* | YjbJ** | YjbJ* | RplT** | RplT* | RpmE* | phylogroup | species |
|---|---|---|---|---|---|---|---|---|---|
| SB5404 | 3850.892 | 7701.426 | 4152.821 | 8305.38 | 4768.43 | 9535.509 | 7739.169 | Kp1 | *K. pneumoniae* |
| SB5404 | 1 | 1 | 1 | 1 | 1 | 1 | 1 | Kp1 | *K. pneumoniae* |
