## Supplementary figures and images for "*Klebsiella* MALDI TypeR: a web-based tool for *Klebsiella* identification based on MALDI-TOF mass spectrometry"

### Supplemental Figure S3

#### C – Visual exploration

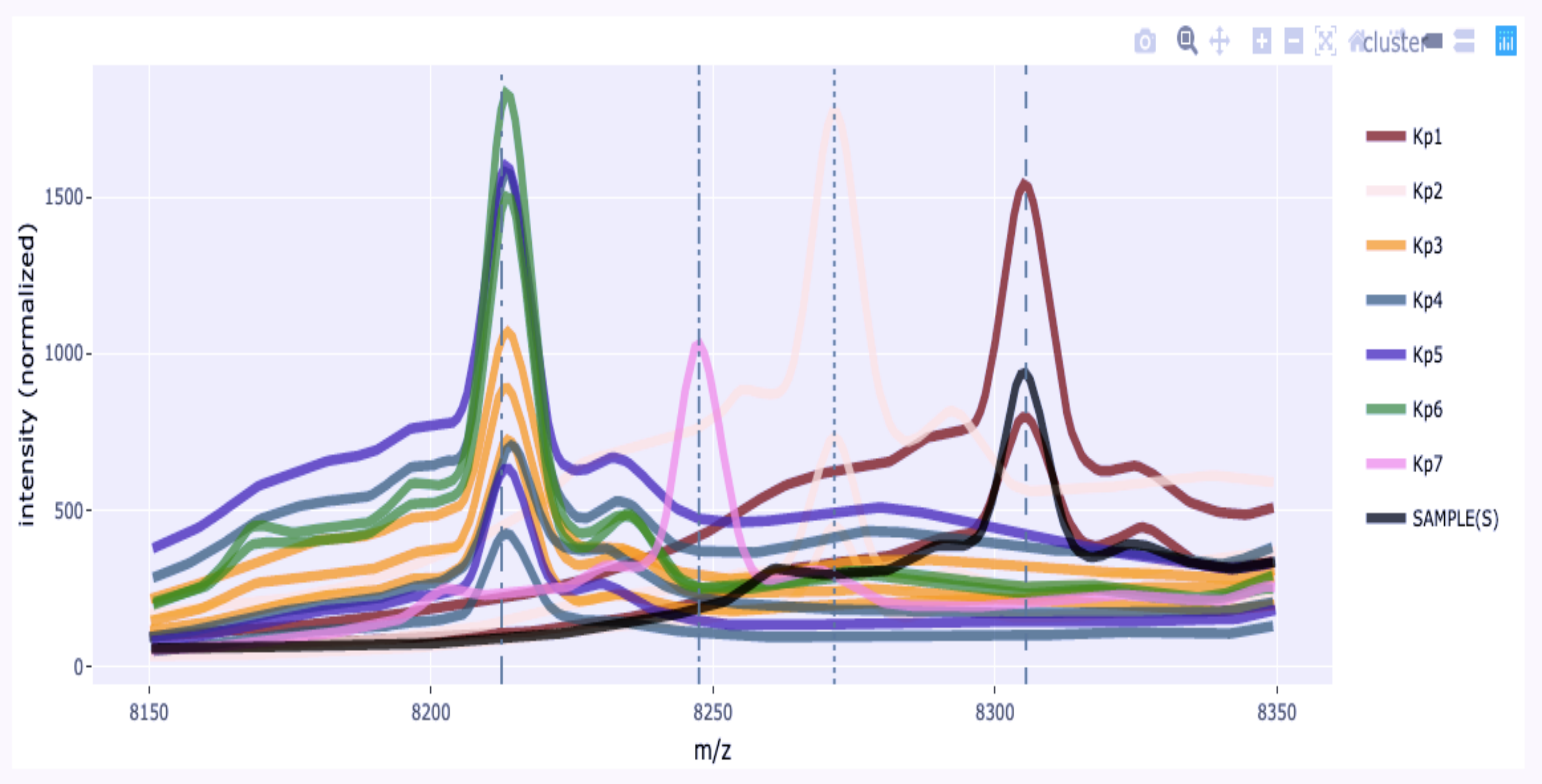
